# Molecular and cellular pathology of monogenic Alzheimer’s disease at single cell resolution

**DOI:** 10.1101/2020.07.14.202317

**Authors:** Federica Marinaro, Moritz Haneklaus, Zhechun Zhang, Alessio Strano, Lewis Evans, Louis-François Handfield, Natalie S. Ryan, Nick C. Fox, Martin Hemberg, Sharad Ramanathan, Frederick J. Livesey

**Affiliations:** Department of Cancer and Developmental Biology & Zayed Centre for Research into Rare Disease in Children, UCL Great Ormond Street Institute of Child Health, 30 Guilford Street, London WC1N 1EH; Wellcome Sanger Institute, Wellcome Genome Campus, Hinxton CB10 1SA, UK; Quantitative Biology Initiative, Harvard University, 52 Oxford Street, Cambridge MA; Department of Stem Cell and Regenerative Biology, Harvard University, Cambridge MA; Dementia Research Centre, Department of Neurodegenerative Disease, UCL Queen Square Institute of Neurology, Queen Square, London; UK Dementia Research Institute at University College London, London, UK

## Abstract

Cell and molecular biology analyses of sporadic Alzheimer’s disease brain are confounded by clinical variability, ageing and genetic heterogeneity. Therefore, we used single-nucleus RNA sequencing to characterize cell composition and gene expression in the cerebral cortex in early-onset, monogenic Alzheimer’s disease. Constructing a cellular atlas of frontal cortex from 8 monogenic AD individuals and 8 matched controls, provided insights into which neurons degenerate in AD and responses of different cell types to AD at the cellular and systems level. Such responses are a combination of positively adaptive and deleterious changes, including large-scale changes in synaptic transmission and marked metabolic reprogramming in neurons. The nature and scale of the transcriptional changes in AD emphasizes the global impact of the disease across all brain cell types.

**One Sentence Summary:** Alzheimer’s disease brain atlas provides insights into disease mechanisms

## Main Text

While many genetic and non-genetic Alzheimer’s disease risk factors have been identified, a unified understanding of cellular and molecular pathogenesis in the disease is lacking (*1*). Monogenic AD provides a defined genetic starting point for studies of disease pathogenesis, circumventing the variation introduced by the genetic heterogeneity (*2*, *3*) and co-morbidities of sporadic, late-onset disease (*4*, *5*). Mutations in amyloid protein precursor (*APP*) and presenilin-1 (*PSEN1*) genes cause autosomal dominant forms of early-onset Alzheimer disease (*5*–*8*). These genes encode for a protease and one of its substrates, and act in a common pathogenic pathway (*9*). Given the early onset of monogenic AD (*4*, *5*), comparisons with age-matched controls reduces the impact of age as a confounding factor for interpreting cellular changes (*10*–*12*).

We analyzed gene expression in single nuclei from *post-mortem* frontal cortex (Brodmann area 9) of 8 individuals with monogenic AD carrying *PSEN1* Intron4, M146I or *APP* V717 mutations, and 8 age- and gender-matched controls (Fig. 1A, Data S1). Neuronal and non-neuronal/glial nuclei were separated by FACS, enabling equal representation of neurons and glial cells in the dataset (NeuN^+^ and NeuN^−^ respectively; Fig. S1, A to D). Droplet-based single-nucleus RNA sequencing (snRNA-seq; see Methods for details) was carried out separately for neuronal and glial nuclei. A two-step process using cell types of human middle temporal gyrus from snRNA-seq data generated by the Allen Institute for Brain Science as a reference (*13*) resulted in a final dataset of 89,325 high confidence nuclei (64,408 from controls and 24,917 from monogenic AD), which was used for all subsequent analyses. Consistent with previous studies of sporadic (*8*) and monogenic AD (*4*, *5*), nuclear sorting identified a marked reduction in the number of neurons (NeuN^+^) in monogenic AD patients with *PSEN*1 or *APP* mutations (Fig. 1B; Fig. S1, B and C) (*14*, *15*). The neuronal nuclei from non-demented control brains had a higher mRNA content compared to glial nuclei (*13*, *16*) (Fig. 1C and Fig. S1, E and F). In contrast we found that there was a marked reduction in the mRNA content of neuronal and glial nuclei from monogenic AD cortex, compared with their counterparts in non-demented controls (Fig. 1C and Fig. S1, E and F).

**Fig. 1.**
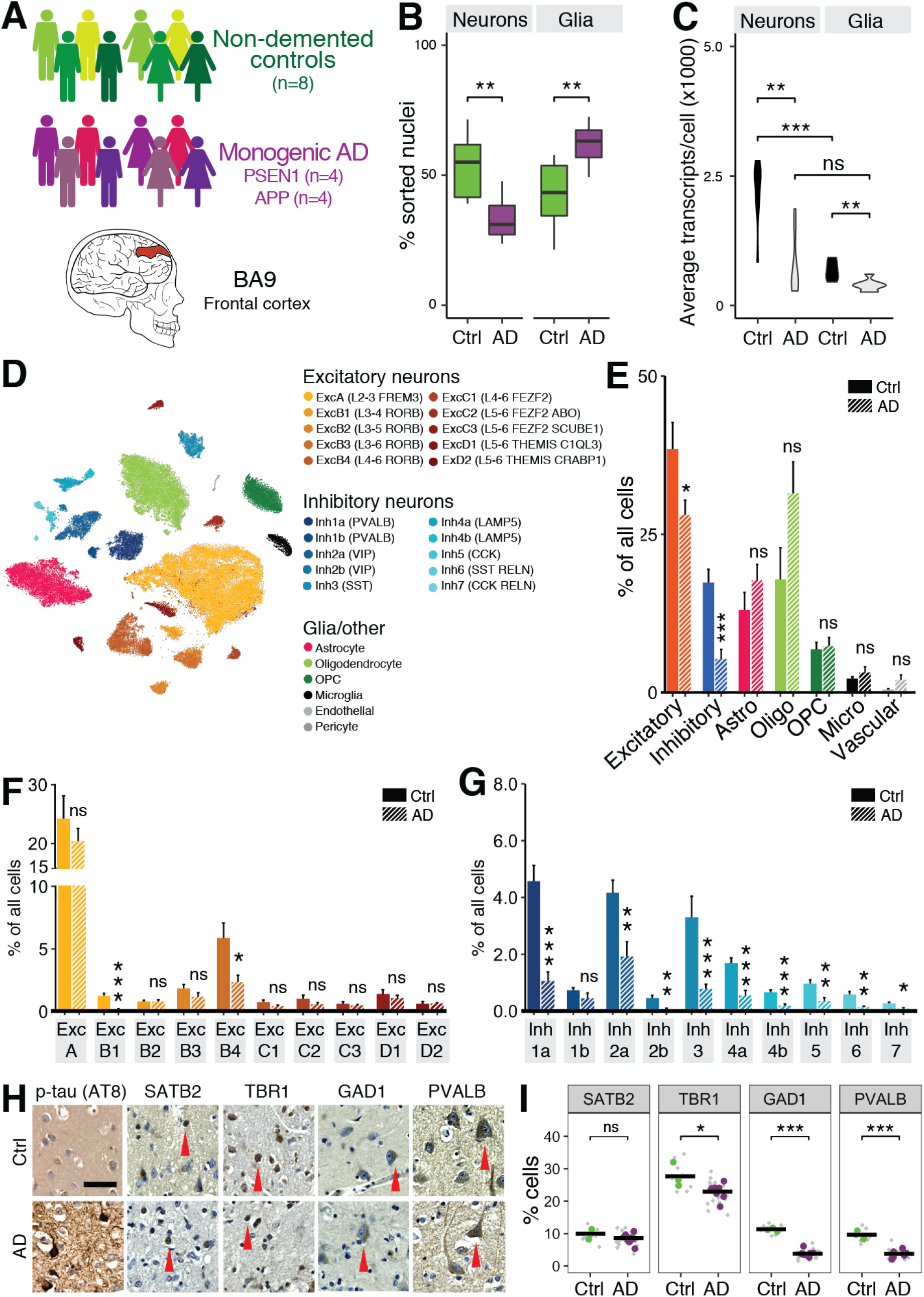
Degeneration of both excitatory and inhibitory neurons in the monogenic AD cortex. (**A**) Frontal cortex (Brodmann area 9, BA9) was analyzed by snRNAseq in 8 monogenic AD patients and 8 matched non-demented controls. (**B**) Comparison of number of neurons (NeuN^+^) and glia (NeuN^−^) in control and AD cortex detected by FACS (\*\**P* < 0.01, two-sided Student’s t-test). (**C**) Comparison of mRNA content per nucleus in control and AD. (\*\**P* < 0.01, \*\*\**P* < 0.001; ns, not significant; Bonferroni-corrected two-sided Student’s t-test). (**D**) Mapping of cell types (t-SNE projection of 89,325 nuclei). Cell types were annotated by similarity to the Allen Institute cell type atlas. (**E-G**) Quantification of normalized numbers of major cell types (E), excitatory (F) and inhibitory neuronal subtypes (G) in control and AD cortex (Mean ±SEM; \**P* < 0.05, \*\**P* < 0.01, \*\*\**P* < 0.001; ns, not significant; two-sided Student’s t-test). (**H,I**) Immunostaining for phosphorylated tau and major classes of excitatory (TBR1, SATB2) and inhibitory neurons (GAD1, PVALB) in the *post-mortem* cortex (H; *arrowheads,* examples of neurons expressing each protein; *scale bar*, 100 μm) and quantification (I; counts per 500 μm unit width of cortex). Three sections spanning the entire cortical thickness (diamonds) were quantified per individual and then averaged (dots); (\**P* < 0.05, \*\*\**P* < 0.001; ns, not significant; two-sided Student’s t-test).

Previous studies of sporadic AD brain identified loss of excitatory neurons from the cerebral cortex, most notably from entorhinal cortex early in the disease (*17*–*20*). To determine the degree of selective neuronal vulnerability in monogenic AD, we compared the proportions of the 10 types of excitatory and 10 types of inhibitory neurons, and 6 glial cell types in the monogenic AD frontal cortex with that of non-demented controls (Fig. 1D-G). For broad categories of neuronal types, we found that both excitatory and inhibitory neurons were significantly reduced in AD (Fig. 1E). In contrast, the proportions of glial cells were not significantly different, although there was a relative increase in both astrocytes and oligodendrocytes, most probably reflecting the loss of neurons (Fig. 1E). While excitatory neurons were broadly lost from the AD cortex, certain classes of excitatory neurons were disproportionately reduced, most notably classes of layer 3/4 (ExcB1) and layer 4-6 neurons (ExcB4; Fig. 1F). In contrast, almost all subtypes of interneurons were significantly reduced in the frontal cortex of monogenic AD patients (Fig. 1G). These findings were confirmed by cell counting in tissue sections from the same monogenic AD and control individuals (Fig. 1, H and I), with, for example, one of the most abundant interneuron subtypes (Parvalbumin^+^, PVALB) reduced by 62% in AD cortex (Fig. 1I). The large-scale loss of inhibition due to interneuron degeneration would be expected to impair excitation:inhibition balance, leading to epilepsy. This is consistent with the high incidence of seizures observed in monogenic AD (*4*, *5*, *21*–*24*), with five of the AD patients studied here manifesting seizures and/or myoclonus, and with the hypothesis that seizures may occur before widespread neurodegeneration and even before clinical symptoms of dementia (*25*).

For the analysis of gene expression changes and their relevance to AD pathogenesis, we focused on those alterations shared between *PSEN1* and *APP* AD. To interrogate the biological relevance of changes in gene expression in each cell type, we analyzed not only individual genes exhibiting the strongest differences in gene expression in each cell type between monogenic AD and non-demented controls, but also functional groups (Fig. 2A and 3, A to C; Fig. S5 to S8; Data S2 and S3). Down-regulation of gene expression dominates in many cell types, including excitatory neurons, inhibitory neurons, oligodendrocytes and oligodendrocyte precursor cells (Fig. S5, A, C and E). Notably amongst functional categories, many genes encoding pre- and post-synaptic proteins involved in synaptic transmission were down-regulated in both excitatory and inhibitory neurons (Fig. 2A; Fig. S6B). Furthermore, both inhibitory and excitatory neurotransmitter receptors were downregulated in expression, as were genes required for GABA production in interneurons. Therefore, in addition to neuron loss, reduced expression of synapse and neurotransmission genes may contribute to declining neurological function and to the development of epilepsy in monogenic AD.

**Fig. 2.**
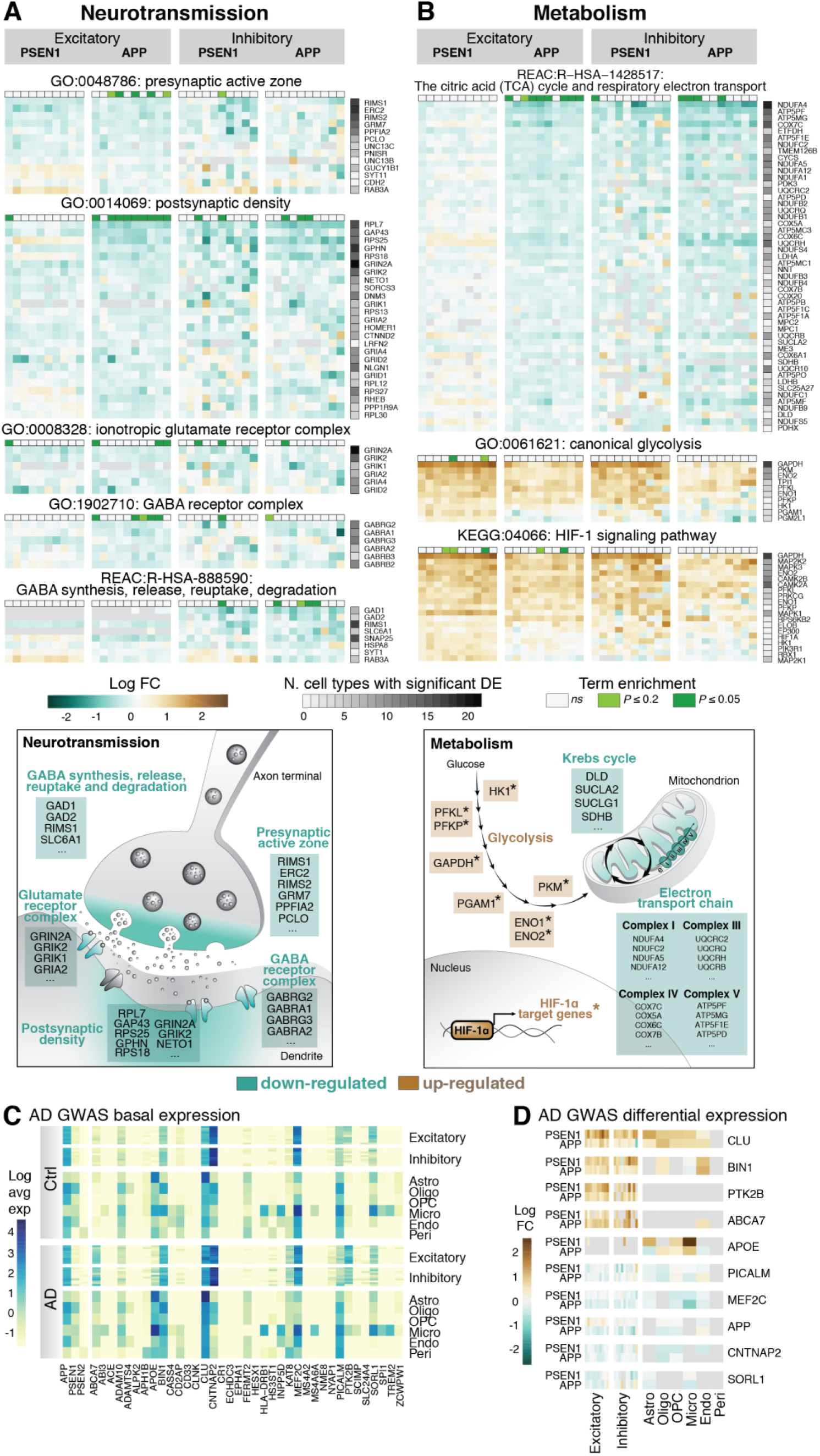
Differential gene expression in AD neurons. (**A**) AD neurons downregulate gene expression networks and pathways required for both excitatory and inhibitory neurotransmission. (**B**) Downregulation of elements of the citric acid cycle and large numbers of genes required for the electron transport chain in AD neurons is accompanied by upregulation of glycolysis genes and the HIF-1 pathway, indicative of metabolic reprogramming. Heatmaps represent log-transformed fold differences in gene expression between APP or PSEN1 AD and matched controls. Term enrichment (top) and the number comparisons in which genes are significantly changed (heatmap, right) are shown. Diagram below summarizes the net effect on neurotransmission. (**C**) Genes identified by GWAS as contributing to risk of developing late-onset Alzheimer’s disease are expressed in many different cell types in both non-demented individuals and monogenic AD. Log-transformed average gene expression per cell type in nuclei from non-demented controls (upper) or AD cases (lower) are shown. (**D**) Many AD GWAS genes that have altered expression in AD, compared with non-demented controls, do so in neurons. Both up- and down-regulation of AD GWAS genes is observed. Cell type-specific differential expression of AD GWAS genes in AD brains (log-transformed fold change) is shown.

**Fig. 3.**
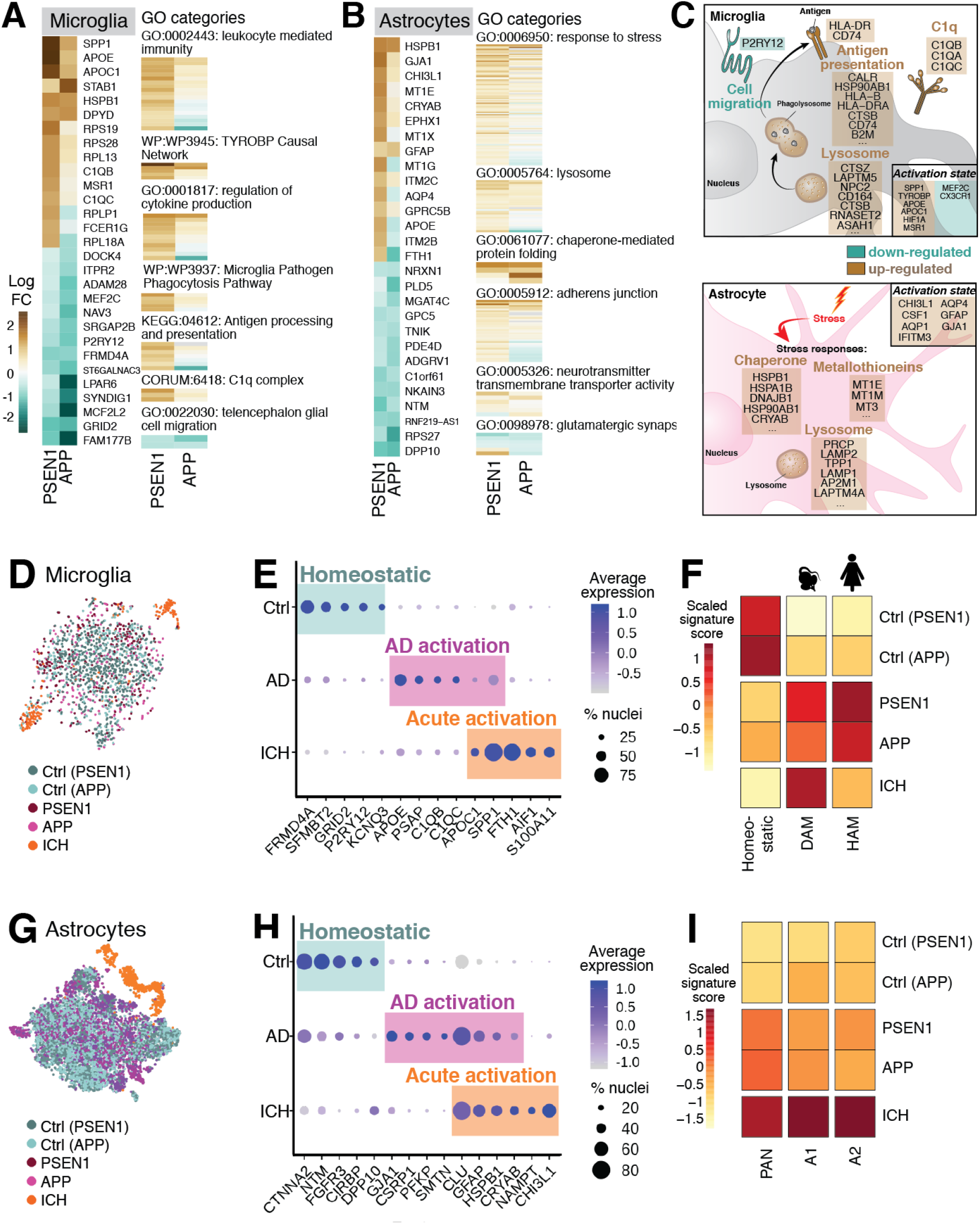
AD-specific phenotypic changes in microglia and astrocytes. (**A-C**) Microglia and astrocytes display complex phenotypic changes in AD. For each cell type, genes with the most significant changes in expression are shown, together with notable functional categories enriched in each cell type. Log fold-change in expression are shown. Key functional changes in AD in each cell type are summarized in cartoons in (C). (**D**) Microglia in AD brain display a disease-specific phenotype, distinct form controls and acute activation due to intracerebral hemorrhage, as illustrated by distribution of disease groups in two-dimensional (t-SNE) projection. (**E**) Expression of genes defining different molecular states of microglia in controls, monogenic AD and intracerebral hemorrhage (ICH). *Dot size*, percentage of nuclei with non-zero expression; *colors*, scaled average expression. (**F**) The AD microglial gene expression signature is more similar to that of human Alzheimer’s microglia (HAM) than murine damage-associated microglia (DAM). (**G-I**) Astrocytes similarly demonstrate an AD-specific activated phenotype, distinct from acute activation. Elements of previously described reactive astrocyte phenotypes (Pan-reactive and A1-/A2-specific) are found in AD astrocytes, but to a lesser degree than in acute activation due to brain hemorrhage (ICH).

In addition to defects in neurotransmission, differential gene expression pointed to a marked switch in neuronal metabolism. A large number of genes encoding multiple elements of the mitochondrial electron transport chain, as well as a number of enzymes required for the Krebs cycle, were downregulated in monogenic AD neurons (Fig. 2B and Fig. S6B). This was seen across multiple neuronal types and indicates widespread mitochondrial dysfunction and defects in oxidative phosphorylation in neurons. Accompanying this was an upregulation of genes involved in glycolysis, many of which are known HIF-1 targets (Fig 2B, Fig S6B and Data S3). Reduced expression of electron transport chain complexes and changes in activity of Krebs cycle enzymes have both previously been noted in AD neurons (*26*–*28*), and the upregulation of glycolysis genes is consistent with functional imaging evidence for a relative increase in aerobic glycolysis in sporadic AD (*29*). The combination of both reduced oxidative phosphorylation and increased glycolysis is reminiscent of metabolic reprogramming observed in cancer (*30*) and during immune cell activation (*31*), which is typically associated with increased metabolic demands. Since mitochondrial dysfunction and oxidative damage are thought to be early events in AD (*32*), it is possible that the shift to glycolysis is required to meet the neurons’ metabolic needs and as a protective mechanism to generate antioxidants. It will be important to determine whether mitochondrial dysfunction or oxidative stress are the primary drivers of metabolic reprogramming in monogenic AD, and more importantly whether this is a protective or pathological process.

To determine possible links between monogenic and sporadic AD, we examined the cellular expression of sporadic AD GWAS-associated genes (*2*, *3*) in monogenic AD patients. Mapping the set of 37 currently known sporadic AD GWAS-associated genes (*2*, *3*), as well as *APP* and *PSEN1/2*, to our dataset, we found that almost all were expressed in at least one cell type in monogenic AD or non-demented controls (Fig. 2C), with many expressed in neurons. Of note, nine AD GWAS genes show significant changes in at least two neuronal subtypes in monogenic AD. This includes genes that are up-regulated (Fig. 2D), despite the overall reduction in gene expression in the monogenic AD cortex (Fig. S5A). For instance, clusterin (CLU), the tau kinase PTK2B (*33*), the APP-processing regulator ABCA7 (*34*) and the regulator of intracellular trafficking BIN1 were all upregulated in neurons (Fig. 2D). Conversely, SORL1, an intracellular sorting receptor for APP (*35*), the endocytosis regulator PICALM, the transmembrane protein CNTNAP2 and the transcription factor MEF2C were down-regulated in neurons (Fig. 2D). The net consequence of these changes is a mixture of protective and pathogenic effects. For example, increasing clusterin levels could potentially be a response to endolysosomal dysfunction (*9*, *36*), and would be predicted to support increased flux through that system. In contrast, reducing SORL1 levels would accelerate pathogenesis, as loss of function *SORL1* mutations are themselves causal for monogenic AD (*37*).

As in neuronal cells, all glial cell types had altered gene expression in monogenic AD (Fig. 3 and S8). Microglia and astrocytes exhibited signs of activation due to inflammatory or damaging stimuli (Fig. 3, A to C; Fig. S8, A and B). Specifically, APOE, SPP1 and complement CQ1 were upregulated in microglia (*38*) and GFAP, CHI3L1 and GJA1 were upregulated in astrocytes (*39*–*41*). In addition, microglia exhibited hallmarks of innate immune cell activation, with upregulation of genes essential for antigen presentation, C1q components, and lysosome components (Fig. 3, A and C; Fig. S8A). Astrocytes also demonstrated several signatures of cellular stress, such as increased expression of molecular chaperones and metallothioneins, and upregulation of lysosomal genes (Fig. 3, B and C; Fig. S8B).

The inflammatory response in monogenic AD was distinct from that in an individual with intracerebral hemorrhage (ICH). For astrocytes and microglia, cells from the ICH cortex formed specific clusters distinct from both monogenic AD and non-demented controls (Fig. 3, D and G). ICH microglia expressed genes associated with acute activation, including SPP1, FTH1 and S100A11 (*42*) (Fig. 3E). The activation of monogenic AD microglia was distinct from that of ICH (Fig. 3E; Data S4). In particular, the monogenic AD microglial phenotype was more similar to the recently described human AD microglia phenotype (HAM; Fig. 3F) than to the murine damage-associated microglia phenotype (DAM; Fig. 3F) (*43*). In contrast, ICH microglia were more similar to murine DAM than human AD microglia (Fig. 3F). A similar trend is observed in AD astrocytes, which have increased expression of a number of genes that have been identified in reactive astrocytes in mouse models of AD (Fig. 3, G to I; Data S4) (*39*). In contrast, reactive astrocyte genes were expressed at a higher level in ICH astrocytes (Fig. 3I). Overall, we conclude that microglia and astrocytes in the monogenic AD brain have an activation phenotype that is disease-specific and distinct from acute activation due to brain hemorrhage.

To complement the analysis above that focused on intracellular signaling, we also studied how changes in gene expression affect intercellular signaling in monogenic AD. To do so, we analyzed co-expression of ligands and their cognate receptors across different cell types, comparing AD and non-demented controls. This analysis revealed an overall decrease in potential cell-cell signaling among neurons in monogenic AD (Fig. 4A; Fig. S9, A to D), but an increase in neuron-microglia and neuron-astrocyte signaling (Fig. 4A; Fig. S9, A to D). Some changes in neuron-glia signaling are likely to be positively adaptive to the ongoing disease process and others deleterious (Fig. 4, B to D, Fig. S9, E to H). These include a positive adaptive change in neuronal scavenging of granulins, with both upregulation of GRN expression by microglia and increased neuronal expression of the SORT1 receptor (Fig. 4B). These changes would increase neuronal accumulation of granulins, improving lysosomal function in *PSEN1* and *APP* mutant neurons, which is compromised by these mutations (*9*). Conversely, microglial homeostasis via the chemokine CX3CL1 appears compromised by reduced expression in neurons of both the ADAM chemokine processing enzymes (*44*) and the CX3CR1 receptor (Fig. 4C). Similarly, neuron-oligodendrocyte/astrocyte signaling via neuregulins is also compromised in AD, with down-regulation of neuronal neuregulin expression and reduced expression of the relevant receptor ERBB4 (*45*) in OPCs and oligodendrocytes and EGFR in astrocytes (*46*) (Fig. 4D).

**Fig. 4.**
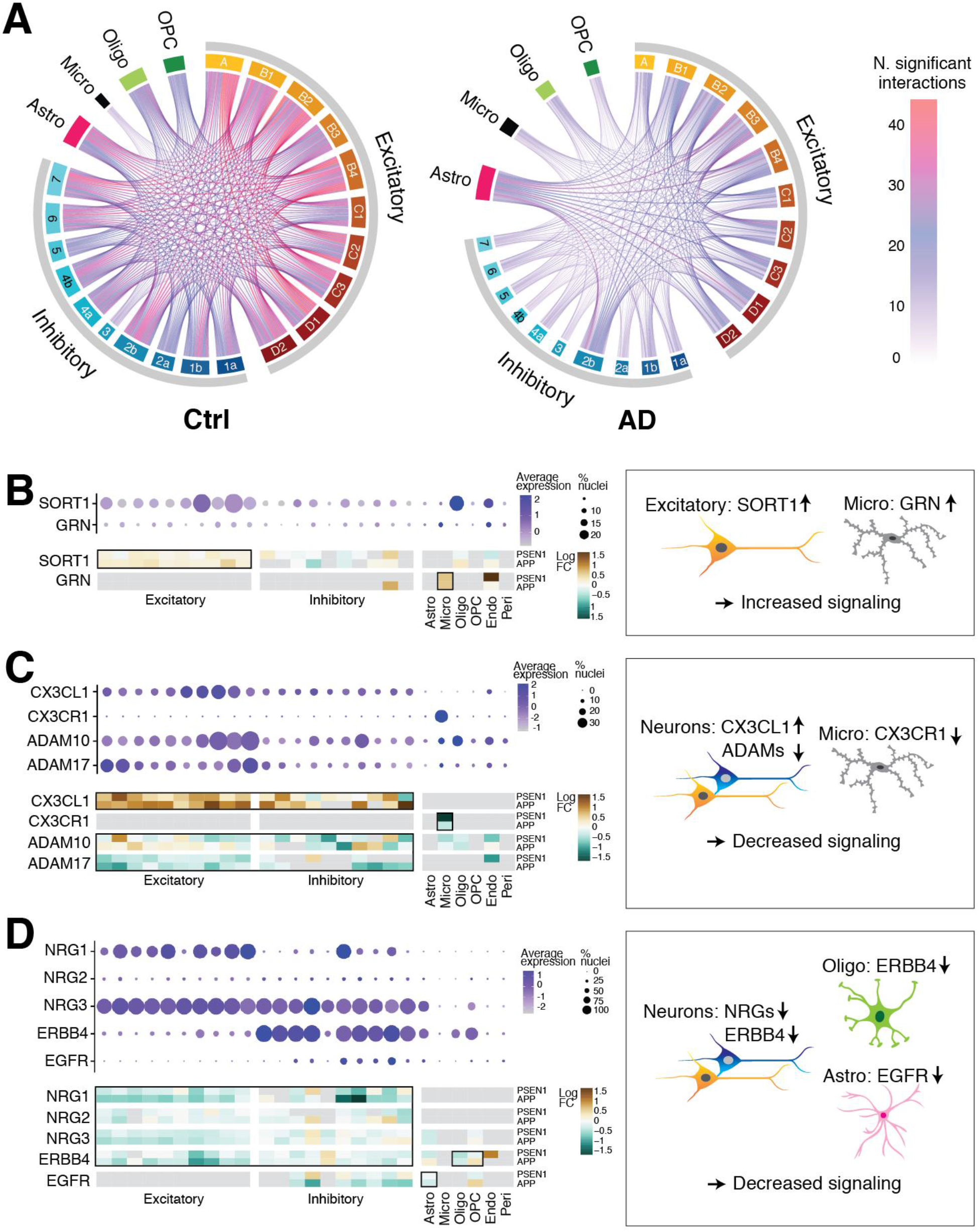
Changes in neuron-glia crosstalk in monogenic AD are a combination of positively adaptive and deleterious responses. (**A**) Potential cell-cell interactions are reduced in AD, whereas as interactions involving either microglia or astrocytes are increased in AD. Circos plots depicting the number of significant potential intercellular molecular interactions in control (left) and monogenic AD (right). (**B-D**) Examples of positively adaptive (B) and deleterious (C-D) changes in receptor-ligand interactions between neurons and glia. Basal expression of relevant genes is shown as dot plots (*Dot size*, percentage of nuclei with non-zero expression; *colors*, scaled average expression). Heatmaps show log-transformed fold change between APP and PSEN1 and matched controls. Differential expression in specific cell types of particular biological relevance is highlighted by black boxes and the likely biological net effect summarized diagrammatically.

Single-cell analysis of the frontal cortex in monogenic AD led to a number of insights into the molecular and cellular pathology of monogenic AD, conserved between patients carrying the *PSEN1* and *APP* mutations. First, we observed widespread degeneration of almost all classes of inhibitory interneurons in monogenic AD, consistent with high incidence of epilepsy in the patients. Second, we found that neuronal cells undergo metabolic reprogramming, similar to cancer cells. Third, we observed both adaptive and deleterious changes in neuronal and glial cells in monogenic AD patients. The presence of positive adaptive changes may explain the relatively slow development of clinical symptoms over decades (*9*, *47*–*51*), and may also point towards resilience mechanisms that support survival of these neurons late in the disease process. In addition to specific gene expression changes, the pervasive and global nature of the disease-associated changes across multiple cell types, and almost all cells within each class, underlines the global nature of the disease in its latter stages. As such, it is consistent with a disease process that begins in early adulthood (*52*), and supports the hypothesis that successful treatments for monogenic AD will likely need to be administered decades before the typical age at onset of clinical symptoms (*4*, *5*).

## Acknowledgments

We thank Rebecca Hodge (Allen Institute) for technical advice and the Cellular Genetics Informatics team at the Wellcome Sanger Institute.

## Funding

Research in this report was supported by Wellcome (Investigator Award to F.J.L.), Alzheimer’s Research UK (Stem Cell Research Centre), Dementias Platform UK (Stem Cell Network), Great Ormond Street Hospital Charity (Stem Cell Professorship, F.J.L.) and NIMH (R01HD100036, S.R.).

## Author contributions

F.M.: Conceptualization, data curation, investigation, project administration, resources, validation, visualization, writing (original draft); Mo.H.: conceptualization, data curation, formal analysis, software, visualization, writing (original draft); Z.Z.; data curation, formal analysis, software, writing (review and editing); A.S.: data curation, formal analysis, software, visualization, writing (review and editing); L.F.H.: data curation, formal analysis, software; N.S.R. resources, writing (review and editing); N.C.F.: supervision, writing (review and editing); Ma.H.: supervision, writing (review and editing); S.R.: supervision, writing (review and editing); F.J.L.: conceptualization, funding acquisition, project administration, supervision, writing (original draft).

## Competing interests

Authors declare no competing interests.

## Data and materials availability

Raw snRNA-seq data and processed count matrix will be uploaded to ArrayExpress at the EBI. Code is available from the corresponding authors upon reasonable request.

## Supplementary Materials

Materials and Methods

Figures S1-S9

Data S1

## Notes

### Competing Interest Statement

The authors have declared no competing interest.

