## Supplementary materials for "Molecular and cellular pathology of monogenic Alzheimer’s disease at single cell resolution"

**This PDF file includes:**

Materials and Methods
Figs. S1 to S9
Data S1

### Materials and Methods

Cases

Frozen *post-mortem* cerebral cortex samples from monogenic Alzheimer's disease (AD) and non-demented control cases were obtained from Queen Square Brain Bank (QSBB), UCL Institute of Neurology, London, UK and from the Sudden Death Brain Bank, University of Edinburgh. The use of human *post-mortem* tissues for this study has been approved with Research Ethics Committee reference ID 08/H0718/54+5 for the cohort from QSBB and reference ID 16/ES/0084 for the cohort from the University of Edinburgh. All AD cases had been genetically tested and this study included three cases carrying a PSEN1 intron 4 mutation, one case carrying PSEN1 M146I mutation and four cases carrying APP V717 mutations. All AD cases met the neuropathological CERAD criteria for definitive AD, with neurofibrillary pathology spread to the occipital cortex reaching Braak and Braak stage 6 and Aβ pathology in the cerebellum reaching Thal stage 5. Neurologically normal controls were cases that had died without history of dementia or neurological disease, with *post-mortem* brain showing absence of any sign of neurodegeneration. Detailed patient data are provided in Data S1. This study was approved by the QSBB and the University of Edinburgh tissue request committees. The cohort in this study is composed by a total of 8 AD cases and 8 non-demented controls (years, control: 45-66 vs. AD:41-66), half males and females. AD and control individuals were matched by sex, age of death, and *post-mortem* delay.

Processing human post-mortem brain samples for single nucleus sequencing

For this study, analysis focused on the frontal cortex, Brodmann area 9 (BA9). Cortical samples were prepared by cryostat sectioning of the tissues in order to encompass the entire cortical thickness. White matter and meninges were dissected from cortical grey matter prior to tissue processing. Matched control and AD samples were processed for isolation of nuclei in parallel to minimize potential batch effects. Nucleus isolation was performed as previously described (*13*), with minor changes. 150-300 mg of sectioned brain tissue was minced in cold, pH 8.0, RNase-free homogenization buffer (10mM Tris buffer, 250 mM sucrose, 25 mM KCl, 5 mM $MgCl_2$, 0.1 mM DTT, 1X protease inhibitor cocktail, 1% RNasin Plus RNase inhibitor, 0.1% Triton X-100 in nuclease-free water) before homogenization in a glass Dounce on ice. Homogenate was filtered through a 30 μm cell strainer (MACS SmartStrainers, Miltenyi Biotech), before recovery of nuclei by centrifugation.

Recovered nuclei were counted using a hemocytometer, diluted to 830 nuclei/μl and split into tubes for subsequent staining with fluorophore-conjugated anti-NeuN antibody (Millipore, FCMAB317PE; 4μl/1x$10^6$ nuclei) or mouse IgG1k Isotype control (Millipore, FCMAB230P; 4μl/1x$10^6$ nuclei). Prior to FACS, nuclear DNA was labelled with DAPI at a final concentration of 1μg/ml. Single nucleus sorting was carried out on a Sony SH800Z cell sorter. Single nuclei were captured by gating on $DAPI^+$ events, excluding cell debris and duplets, and subsequently on NeuN expression. Sorted nuclei were processed by single-nucleus capture using the Chromium™ Single Cell 3' Reagent Kit (v2 Chemistry; 10X Genomics). The 10X protocol for capture and library preparation was used without modification. Sorted nuclei from matched control and AD samples were processed in parallel on the same 10X chip to minimize batch effects. Single-nucleus libraries from individual samples were pooled and sequenced using a HiSeq 4000 Sequencing System (Illumina).

Data processing

Gene expression count matrices were generated from Fastq files using Cell Ranger 3.0.0 (10X Genomics). In order to increase transcriptional information in nuclei, intronic regions were included in the reference transcriptome(*53*). This was achieved by reclassifying intronic regions in the GTF file (release 93 from Ensembl; Homo_sapiens.GRCh38.93.gtf) to be processed as 'exons' by 'Cellranger mkref'. This approach was similar but not identical to the suggested workflow by 10X Genomics to generate a pre-mRNA reference. We first included the complete genomic region encompassing all exons and introns for each gene as 'exons' in the transcriptome reference. However, we noticed that for a number of genes (e.g. SLC26A3) the vast majority of reads were derived from a small intronic region that was of low complexity, particularly intronic polyT regions, but not any gene-specific surrounding sequences, and therefore most likely arose due to mapping artefacts (data not shown). We therefore further altered the transcriptome reference in such a way that genomic regions that are identified by RepeatMasker as simple repeats, low-complexity or interspersed repeats (Smit AFA, Hubley R, Green P (1996-2010) RepeatMasker Open-3.0 http://www.repeatmasker.org) are excluded during mapping. To do this, we excluded reads for which more than 32bp of the alignment overlap with a genomic range associated with any Repeatmask feature. Pre-computed genomic Repeatmask regions for the hg38 assembly were sourced from the UCSC ftp server. We then included only those intronic sub-ranges as 'exons' in the reference transcriptome that were not identified as low-complexity regions. Using bedtools(*54*), all Repeatmask features were merged into contiguous ranges, including any gap between resulting ranges that was smaller than 32 bp. Every range was then reduced by 66 bp upstream and downstream, and the result was subtracted from every transcript region that would be added as 'exon' in the GTF file. Any 98bp read that fully maps within one of the contiguous ranges produced would then be guaranteed to overlap with at least 32 bp of the transcript that is not associated with any Repeatmask features. Finally, for genes that are not protein coding, the genomic regions for other protein coding genes were subtracted, so that reads were preferentially mapped to the latter category, as multi-mapped reads are discarded during STAR alignment as part of the Cell Ranger pipeline.

Following alignment and count matrix generation, the emptyDrops pipeline from the DropletUtils package (v1.4.3)(*55*) was applied to distinguish nuclear profiles from empty droplets. The knee and inflection points were determined separately for each sample after visually inspecting the output of the barcodeRanks function. The values for the knee point above which all droplets are assigned as non-empty ranged from 164 to 2851 UMI. The values for the inflection point, under which droplets are assigned to be empty, ranged from 30-90 UMI. emptyDrops was then run with sample-specific values for the variables *lower* (knee point) and *retain* (inflection point) and barcodes with less than 100 UMI were ignored. emptyDrops was run with 100,000 iterations in the first instance. If any barcodes remained that did not reject the null hypothesis but were limited by the number of iterations (limited=TRUE), emptyDrops was re-run for 500,000 iterations. Barcodes that were predicted by emptyDrops to differ from empty droplet profiles with an FDR $\leq$ 0.01 were included in the further analysis. Seurat objects for each sequencing run were generated following the standard Seurat v3.0.0[5] pipeline, using the Read10X and CreateSeuratObject functions. For each nucleus, the proportion of mitochondrial reads was determined by dividing the sum of all mitochondrially-encoded genes by the total number of reads.

We removed nuclei with particularly high numbers of reads from mitochondrial genes, which are likely contaminated with or solely contain non-nuclear cellular material. Distributions of mitochondrial read percentage varied among samples (data not shown). We therefore determined outliers in a sample-specific manner by excluding nuclei that exceeded twice the IQR (interquartile range) above the third quartile of mitochondrial percentages for each sample. This left 111,850 nuclei that were used for further processing.

Single-cell datasets were merged using the data integration method in Seurat(*56*). Seurat objects for NeuN$^+$ and NeuN$^-$ samples for each individual were merged, normalized (NormalizeData) and the 2000 most variable features were identified using the FindVariableFeatures function. The objects were then integrated using FindIntegrationAnchors and IntegrateData using 75 dimensions in order to take into account the higher dimensionality of the data. Unless otherwise stated, count matrices were analyzed following standard workflows described in Seurat v3 vignettes. Integrated count matrices were first scaled and the number of UMI per nucleus and the percentage of mitochondrial genes was regressed (ScaleData function). Principal component analysis (PCA) was performed on the scaled, integrated count matrix (RunPCA function) and the first 40 principal components were used for dimensionality reduction (RunTSNE (t-distributed Stochastic Neighbour Embedding; t-SNE) as well as a shared nearest neighbour (SNN) modularity optimization-based clustering algorithm(*57*) embedded in Seurat v3 (FindNeighbors and FindClusters functions). The number of principal components to use for dimensionality reduction and clustering was estimated using the ElbowPlot function. FindClusters was run with 100 random starts and a maximum of 100 iterations per random start (n.start=100, n.iter=100). RunTSNE was run with the FFT-accelerated Interpolation-based t-SNE (FIt-SNE)(*58*).

Nearest neighbors per nucleus were determined using the nn2 function of the RANN R package v 2.6.1, which utilizes Arya and Mount's Approximate Nearest Neighbours (ANN) C++ library(*59*). Unless stated otherwise, embeddings of the first 40 principal components were used as input.

Cell type annotations were transferred from reference datasets to the query using the label transfer feature of Seurat v3 (FindTransferAnchors and TransferData) based on the first 30 dimensions, unless otherwise stated.

In-house frontal cortex atlas generation

To annotate cell types in our dataset in a robust and unbiased way and generate an in-house frontal cortex atlas, we used the well-annotated dataset of cell types in the human middle temporal gyrus (MTG) that was generated by the Allen Institute for Brain Science as a reference(*13*). We only included high quality samples of healthy controls in order to determine the largest number of cell types that could be captured by snRNA-seq. We chose to exclude one control sample that had particularly low UMI counts per nucleus (CTRL1) and the donor that died from a cerebral hemorrhage (CTRL6). We then used the workflow described in "Data processing" to merge the remaining six samples and process the data.

We next identified the broad cell classes that make up the majority of cells in the human cortex (excitatory and inhibitory neurons as well as astrocytes, oligodendrocytes, OPC, microglia and vascular cells)(*13*) using the label transfer feature of Seurat v3. Since these cell types were well separated by their gene expression profile, we used the following criteria to exclude nuclei of low quality:

- Poorly mapping nuclei (mapping score < 0.6) that also contained a low number of transcripts (450 UMI)
- Nuclei that agreed with the cell type of less than 40 of the 50 nearest neighbors

Finally, the data were clustered with high resolution (res=5) and all clusters that contained a majority of cells that would be excluded by the two criteria above were also excluded from the final reference. In order to define neuronal subclusters, we re-ran the data integration procedure for all excitatory and inhibitory nuclei that were not excluded by the criteria described above. We next mapped the two objects against the excitatory or inhibitory neuronal subtypes of the Allen Institute MTG reference, respectively. For this, we first extracted the excitatory or inhibitory neuronal clusters from the MTG reference and normalized and scaled the data using the standard Seurat pipeline, including regression of the number of UMI. In addition to mapping to this subset of the data, we also mapped the excitatory and inhibitory subsets of the in-house atlas against the complete MTG dataset to identify nuclei that were misclassified in the first instance.

Fine-grain clusters (res=5) were defined and assigned identities based on the neuronal subtype in the MTG dataset to which the majority of nuclei in each cluster were mapped. Clusters that predominantly contained cells mapping to another broad cell type were excluded. For the rest, if more than 50 % of nuclei in a cluster mapped to the same cell type, the Allen Institute MTG cell type identifiers were retained. Otherwise, cell type names of up to the third most common matching cell type were merged. This resulted in 13 excitatory and 25 inhibitory neuron subclusters (data not shown) that could be distinguished. However, not all of these subtypes could be easily identified in the complete dataset including AD samples, as indicated by low mapping scores. We therefore combined several fine cell types into functional groups for which cell clusters with good mapping scores could be identified (data not shown), resulting in 10 excitatory and 10 inhibitory neuronal subtypes. The main identifying genes were retained in the final cell type names, as well as combined layer information for excitatory neurons. Only for 3 interneuron types, identifying genes were changed because they were better distinguished by expression of CCK, SST and RELN (Fig. S4E), these were:

- Inh5 (CCK): containing cells with best matches to "Inh L1-2 GAD1" and "Inh L1-3 PAX6"
- Inh6 (SST RELN): containing cells with best matches to "Inh L1 SST NMBR"
- Inh7 (CCK RELN): containing cells with best matches to "Inh L1-2 PAX6 CDH12"

Layer identity and gene expression corresponded well between clusters of excitatory neurons in our atlas and the MTG dataset, for which cortical layers were physically separated (data not shown) therefore we retained layer information in the final cell type identities. In contrast, inhibitory neurons did not group by layer identity but rather by developmental origin as previously described(*60*) (data not shown) and we therefore only added identifying genes for each cell type. For excitatory neurons, in order to only include nuclei with the most confident identity for the in-house atlas, nuclei that did not agree with the identity of 40 of their 50 nearest neighbors based on 30 principal components were not considered.

Mapping cell types in the monogenic AD atlas

In order to annotate cell type identities for the complete dataset, we first mapped all cells against the internal reference using the steps described above. Similar to the process of cell type identification of the reference, we first mapped broad cell classes (excitatory and inhibitory neurons as well as astrocytes, oligodendrocytes, OPC, microglia and vascular cells). In order to remove nuclei of poor quality, nuclei were excluded that:

- mapped poorly to any of the broad cell types (mapping score < 0.8)
- agreed with the cell type of less than 40 of the 50 nearest neighbors

Data for the remaining nuclei were re-scaled and PCA was repeated.

Excitatory and inhibitory neuronal subtypes were determined separately as described above for the in-house reference atlas. Briefly, excitatory and inhibitory neurons from the complete dataset were extracted and re-integrated separately. They were then mapped against the complete in-house reference or only the subset of excitatory or inhibitory neurons. Cell subsets were clustered (res=0.8) and clusters that contained a majority of non-matching cell types were excluded. In addition, nuclei with low mapping quality (mapping score < 0.8) were excluded. The remaining count matrix was then re-scaled, including regressing out of mitochondrial percentage and number of UMI per nucleus, and PCA, clustering and t-SNE dimensionality reduction re-run to generate the final dataset.

Cell type marker gene identification

Cell type markers were identified based on binary expression in the chosen cell type compared to each other cell type using a formula modified from Hodge et al., 2019(*13*) to identify genes with cell type-specific expression in both non-demented control and monogenic AD samples. For each gene, a specificity score for each cell type ($ß_i$) was calculated separately for the two genotypes (g) (control and AD). The score is based on difference in the proportion of nuclei with non-zero expression of the gene in the cell type of interest ($x_i$) compared to each other cell type ($x_j$). A final specificity score for each gene and cell type was then calculated by adding the control score to the AD score, with a maximum score of 12. The overall formula is:

$$\beta_i = \sum_{g=1}^{g=2} \sum_{j=1}^{j=6} \frac{(x_i - x_j)^2}{x_i - x_j + \varepsilon}$$

where ε is a small constant to avoid division by zero. Markers were identified as genes with highest score in one cell type which also scored less than 0.4 in all other cell types. The threshold was chosen empirically to account for low background expression in other cell types.

Cell type quantification

To calculate changes in cell type composition, we first calculated the proportion of different cell types in each sorted-sequenced sample (2 sorted-sequenced samples ($NeuN^+$ and $NeuN^-$) per brain sample). We then adjusted these proportions by the proportion of $NeuN^+$ and $NeuN^-$ nuclei in each brain sample to obtain adjusted cell type proportions out of all sorted $DAPI^+$ nuclei. Differences in average proportion of cell types in non-demented control and AD brains were tested for significance using a two-sided t-test.

Exclusion of background transcripts

Droplet-based single-cell sequencing techniques contain a significant amount of ambient background RNA, which contribute to the cell-specific count matrix(*61*). Given that nuclei from

AD brains contained lower transcriptional information, background transcripts contribute a larger proportion, which leads to artefacts during differential expression analysis. We therefore excluded 70 genes (Data S2) from functional enrichment analysis and top differentially expressed genes that were particularly high in empty droplets, which are a good approximation for background transcripts. These 70 genes were selected from 311 genes most abundant in empty droplets of NeuN+ control, NeuN+ AD, NeuN- control, and NeuN- AD samples. Such background transcripts can bias differential expression analysis, but the effect is expected to be conserved across different cell types. Therefore, we only excluded 70 genes that show differential expression in more than 75% of NeuN+ or NeuN- cell types.

Differential gene expression

Differential gene expression analysis was performed using the FindMarkers function in Seurat v3 using the normalized raw counts as input with the options logfc.threshold = 0 and min.pct = 0.03. The intracerebral hemorrhage case (CTRL6) was excluded from the analysis and remaining groups that were compared were:

- Control(PSEN1): CTRL5,7,8
- PSEN1: AD5-8
- Control(APP): CTRL1-4
- APP: AD1-4

Genes were considered significant if:

- P value (p_val) < 0.001
- FDR (p_val_adj) < 0.05
- Log fold change (avg_logFC) > 0.25

- Percentage expression in either of the groups (pct.1 | pct.2) > 0.1

Background genes were excluded from significant genes and not included in the generation of Venn diagrams and for functional enrichment analysis. They were also excluded from heatmaps in Figs. 2 and 3, and Fig. S6 and S8, except as part of an enriched GO term or pathway.

Functional enrichment

Differentially expressed genes were used for functional enrichment analysis using g:Profiler(*62*) (R client gprofiler2 v0.1.8). Significantly up- or down-regulated genes in PSEN1 or APP patients for each cell type were separately used as input lists. Enrichment was calculated using the gost function using the additional options:

organism = "hsapiens", exclude_iea = T, evcodes = T, domain_scope = "annotated", correction_method = "gSCS", significant = F

We also tested enrichment for another gene set for each cell type (comparison column):

- 'same_direction': genes that are significant in either APP and PSEN1 comparisons and with the same direction of fold change in the other comparison

Results for all comparisons with $P \leq 0.2$ are listed in Data S3.

Baseline expression of GWAS-linked genes

GWAS-linked genes were extracted from Table 1 in Kunkle *et al.*(*3*) and Table 1 in Jansen et al.(*2*). Baseline expression for each cell type was calculated separately for all non-demented controls (excluding the hemorrhage patient) and monogenic AD patients using the

AverageExpression function in Seurat v3 and heatmaps of log-transformed values were plotted with a lower cut-off of 0.2 were plotted using the R package pheatmap v1.0.12.

<u>Gene selection for differential expression heatmaps</u>

Heatmaps for log-transformed fold change values for indicated genes were plotted using the pheatmap R package. An identical range was chosen that covered all values plotted. The values were plotted for all comparisons in which the gene was detected in at least 5% (all plots except Fig. 2C and Fig. 4, C and D), 8% (Fig. 4, C and D) or 10% (Fig. 2C) of the cells in either of the compared groups.

*Top neuronal genes (Suppl Fig. 6A):*

Genes that were significantly up- or down-regulated in at least one neuronal subtype were ordered by the average fold change in excitatory and inhibitory neuronal subtypes. The 8 strongest up- and down-regulated genes by fold change were included in the plot. For genes specifically up- or down-regulated in excitatory and inhibitory neurons, genes with the 2 largest or smallest difference in the average fold change in excitatory and inhibitory neurons among up- or down-regulated genes were included in the plot.

*Top glial genes (Fig. 3A and B; Fig. S8, C and D):*

For each glial cell type, the 15 most significant genes in the comparison of APP or PSEN1 with the respective matched controls were selected, ordered by average fold change and plotted.

*Enriched pathway genes (Fig. 2A and Fig. 3, A and B; Fig. S6B and S8, A to D):*

Enriched GO terms and pathways that were enriched in up-/down-regulated genes in indicated cell types for either APP or PSEN1 patients were selected based on significance, the number of overlapping genes and biological relevance to the respective cell type. For neurons, we preferentially selected those that showed consistent changes across multiple subtypes. Genes that overlapped with selected terms were extracted from g:Profiler functional enrichment results (Data S3) and plotted. For neurons, overlapping genes were combined for all subtypes. For GWAS-linked genes (Fig. 2C), genes that were significantly differentially expressed in the highest number of cell types were selected and their fold change plotted.

Microglia and astrocyte signature expression

Published gene signatures for homeostatic ('Microglia'), damage-associated ('Neurodegeneration-related') and human AD-associated microglia ('Human-AD-Myeloid' Up; Module U1-3) were extracted from Supplementary Table 2 from Srinivasan *et al.*(*43*). Human gene sets for reactive astrocytes that were originally identified by Liddelow *et al.*(*39*) were extracted from Figure 1c in Diaz-Castro *et al.*(*63*). Expression scores for each signature per cell were calculated using the AddModuleScore function in Seurat. Signature expression in all nuclei of the respective cell types was averaged for each donor. For control and AD groups, mean expression values were further averaged across the values for individuals in each group. Average expression values were plotted using pheatmap.

Glial subclusters

The complete dataset was used to identified subsets of nuclei defined as microglia, astrocytes, OPCs or oligodendrocytes with a proportion of mitochondrial reads below 0.15. Nuclei subsets were analyzed with Seurat v3 and sctransform v0.2.0(*64*) following the vignette "Using sctransform in Seurat" (https://satijalab.org/seurat/v3.0/sctransform_vignette.html). Briefly, raw count matrices were pre-processed using the Seurat function SCTransform, including regression of the number of UMI per nucleus, mitochondrial read percentage, individual of origin as well as NeuN sorting to avoid clustering by individual. Dimensionality reduction of the normalized count matrix by PCA and t-SNE embedding as well as identification of clusters was performed as described above. The number of principal components used for t-SNE embedding and cluster identification was approximated using the ElbowPlot function (microglia: 10, astrocytes: 10, oligodendrocytes: 15, OPC: 20). For astrocytes, small numbers of contaminating neurons and oligodendrocytes were removed after clustering with res=2.5 and pre-processing was repeated. Genes specifically expressed in non-demented controls, monogenic AD patients or an individual who suffered a brain hemorrhage (CTRL6) were determined using the FindMarkers function in Seurat using the normalized raw data as input using the default Wilcoxon Rank Sum test. Gene expression of selected genes was plotted using the DotPlot function and t-SNE plots were generated using the DimPlot function. Genes specific for AD or ICH microglia and astrocytes are summarized in Data S4.

CellPhoneDB cell-cell interaction analysis

Co-expression of ligand-receptor pairs between different cell types were analyzed using CellPhoneDB v2.0(*65, 66*) (https://github.com/Teichlab/cellphonedb).

The dataset was sub-sampled to 80 cells per cell type per genotype. In order to identify glial-neuronal interactions that are conserved among neuronal subtypes, log co-expression and significance values were averaged over excitatory or inhibitory subtypes. The number of significant interactions were visualized with Circos plots using the R package circlize (v 0.4.8)(*67*).

Histology and cell counting in tissue sections

Eight-micron-thick paraffin tissue sections from the frontal cortex of AD and control cases were provided by Queen Square Brain Bank. Commercially available anti-phospho-tau (AT8, Pierce, MN1020), anti-TBR1 (Abcam, ab31940), anti-SATB2 (Santa Cruz, sc-81376), anti-GAD1 (Chemicon, mab5406) and anti-PVALB (Abcam, ab11427) antibodies were used in this study. DAB-based immunostaining was carried out using standard methods (Vector laboratories, ImmPRESS HRP Universal Antibody, MP-7500-15, ImmPACT DAB Peroxidase HRP Substrate, SK-4105). Nuclear counterstaining was with Meyer's hematoxylin. Brightfield images were acquired with Zeiss Axioscan Z1 slide scanning microscope and analyzed with ImageJ version 2.0.0 (Wayne Rasband, National Institutes of Health, USA). Original images were cropped to prepare figure panels.

Cells expressing each of the antigens/proteins studied were counted through the depth of the cortical wall, and their numbers expressed as a proportion of total cell populations per unit width of cortex, as indicated in figures and legends. Data from each anitbody are the mean of 3 counted fields per individual. Given the size differences between non-demented control and monogenic AD cortices, fields compared covered the same width of ventricular surface (500 μm).

Differences in the average percentage of marker-positive cells (of all cells in the region analyzed) between controls and AD cases were tested for significance using a two-sided t-test. Values for each brain sample were calculated as averages of blinded counts on three separate cortical slices.

Code and data availability

Code will be made available upon reasonable request to the corresponding author.

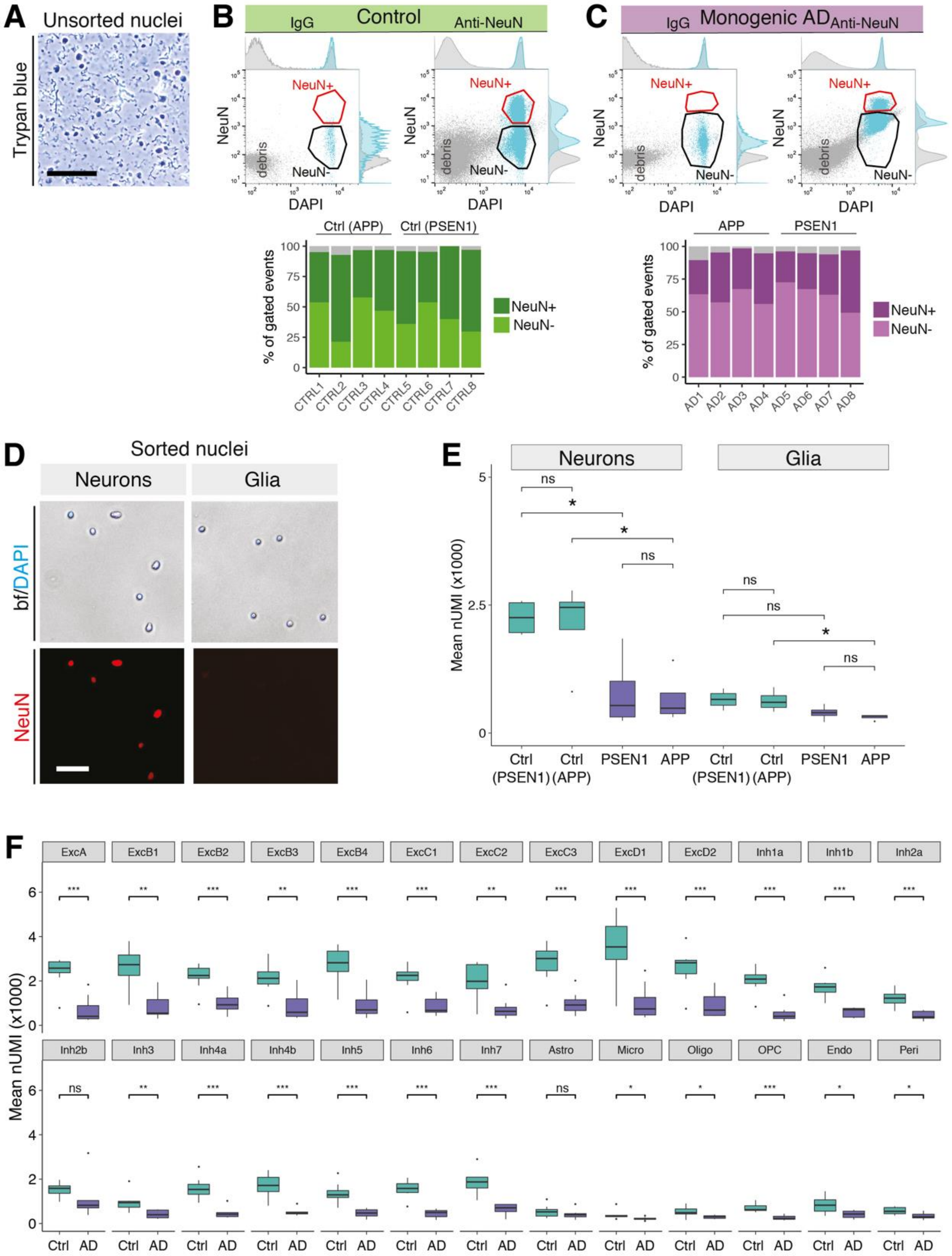


**Supplementary Fig. 1. Human brain nucleus sorting and transcript detection**

(**A**) Unsorted nuclei prepared from frozen post-mortem cerebral cortex stained with Trypan blue (bright field image; scale bar, 100 µm). (**B,C**) Sorting of nuclei from non-demented controls (**B**) and monogenic AD patient cerebral cortex (**C**). Flow cytometry scatter plots of nuclei stained with DAPI (nuclear DNA) and IgG isotype control antibody (left) or stained for DAPI and the neuron-specific nuclear protein NeuN (right). Sorted populations of $DAPI^{+}$/$NeuN^{+}$ and $DAPI^{+}$/$NeuN^{-}$ nuclei are indicated. Bar graphs show the proportion of glial ($NeuN^{-}$) and neuronal ($NeuN^{+}$) nuclei of all $DAPI^{+}$ nuclei for each sample. (**D**) Brightfield (bf) and fluorescence microscopy images of neuronal ($NeuN^{+}$) and glial ($NeuN^{-}$) nuclei after sorting. Red fluorescence represents NeuN staining. (**E**) Comparison of mRNA content per nucleus (average across neurons and glia) in control and AD separated by mutation. (*$P \leq 0.05$; ns, not significant; Bonferroni-corrected two-sided Student's t-test). (**F**) Comparison of mRNA content per nucleus (average across nuclei from each cell type) in control and AD(*$P \leq 0.05$, **$P \leq 0.01$, ***$P \leq 0.001$; ns, not significant; Bonferroni-corrected two-sided Student's t-test).

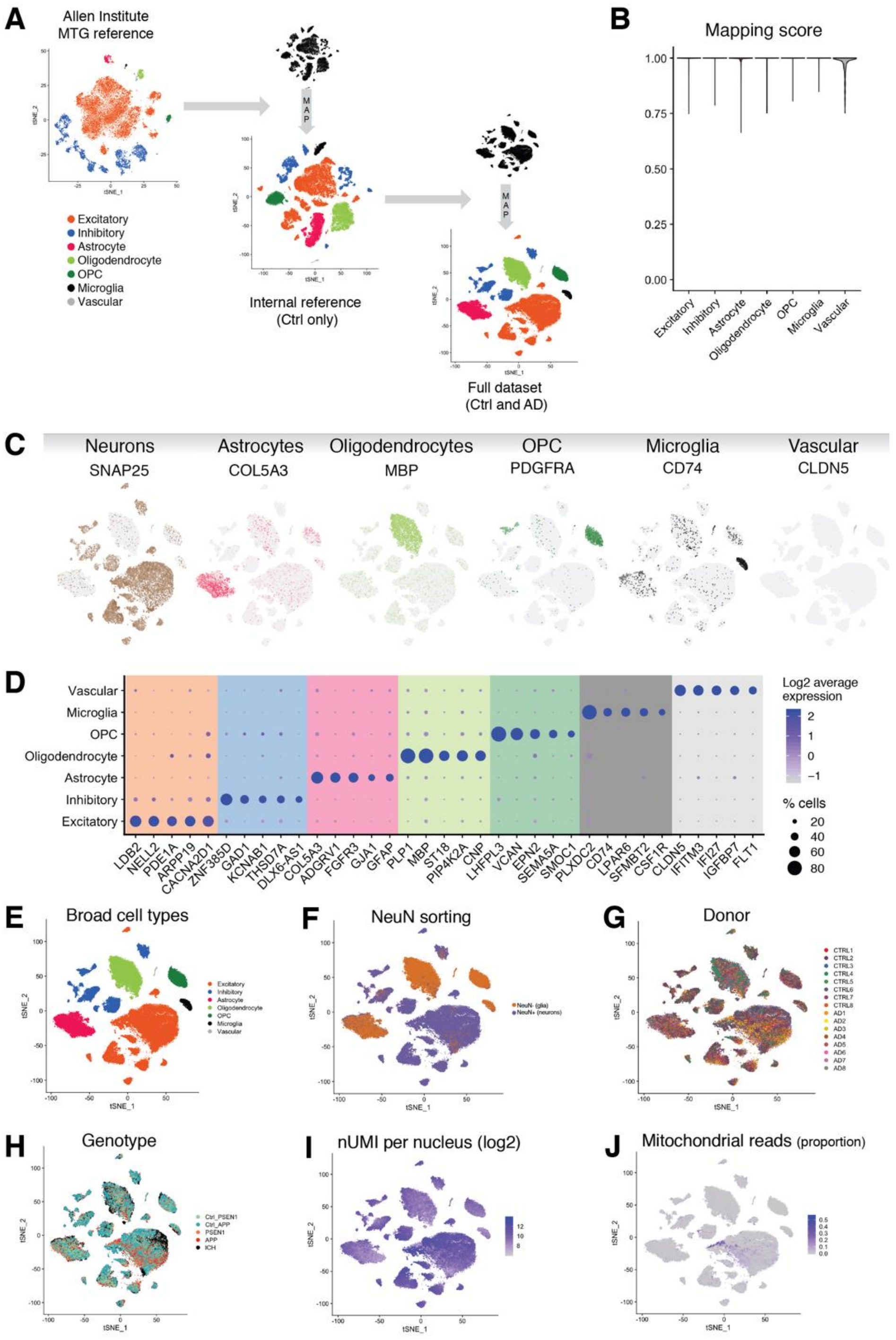


**Supplementary Fig. 2. Mapping strategy and quality control of snRNA-seq data**

(**A-C**) Mapping strategy of snRNA-seq data. (A) Broad cell types of the Allen Institute MTG dataset (*13*) were projected onto an internal reference made up of non-demented control samples. The internal reference in turn was projected onto the complete dataset, including AD samples and controls, to identify broad cell types. t-SNE projections of the three datasets colored by cell type are shown. (B) Violin plot of cell type mapping scores of the complete dataset to the internal reference after cleanup. (C) Expression of marker genes for the indicated cell type overlaid on the t-SNE projection of the complete dataset. (**D**) Cell classes were clearly segmented and categorized by cell-specific/enriched gene expression, as exemplified by expression of 5 genes whose expression is most enriched in each cell type. Size of the dots shows the percentage of nuclei with non-zero expression and color represents the scaled average expression. (**E-I**) Quality control plots of data integration and mapping. t-SNE projection colored by either identified cell type (E), NeuN FACS sorting (F), individual donor (G) and genotype (H) or overlaid with the number of detected transcripts (nUMI; I) and proportion of mitochondrial reads per nucleus (J).

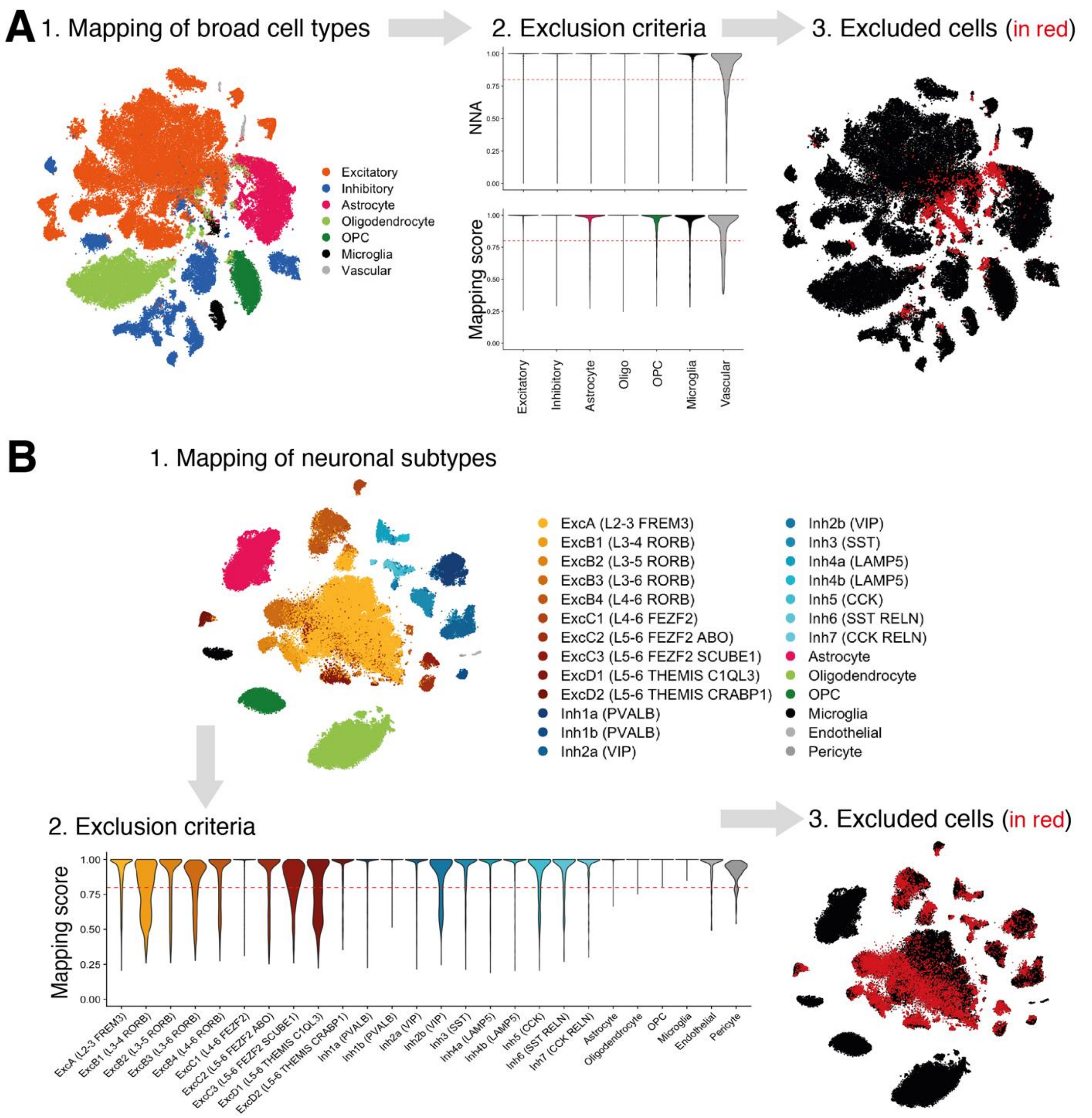


**Supplementary Fig. 3. Dataset clean-up strategy**

(**A**) The initial dataset (111,850 nuclei) was projected onto the internal reference (see Methods for details) to determine broad cell types. Nuclei that did not project with high confidence (prediction score < 0.8) to any of the broad cell types in the reference or whose nearest neighbors did not agree with their predicted identity were excluded (7961 nuclei) and the remaining nuclei taken forward to the next round of data cleanup. t-SNE projection colored by predicted cell type and with excluded cells highlighted are shown. Violin plots show proportion of agreement of cell type prediction with nearest neighbors (upper) and prediction score per nucleus (lower). (**B**) To further remove neurons of uncertain identity, we again projected the remaining 103,889 nuclei to the internal reference (see Methods for details). Excitatory and inhibitory nuclei were projected separately to excitatory and inhibitory reference subtypes, respectively. Nuclei that did not map confidently (prediction score < 0.8) to any cell type (14,564 nuclei) were excluded from the final dataset. Violin plot shows the prediction score per nucleus. Excluded cells are highlighted in a t-SNE projection plot.

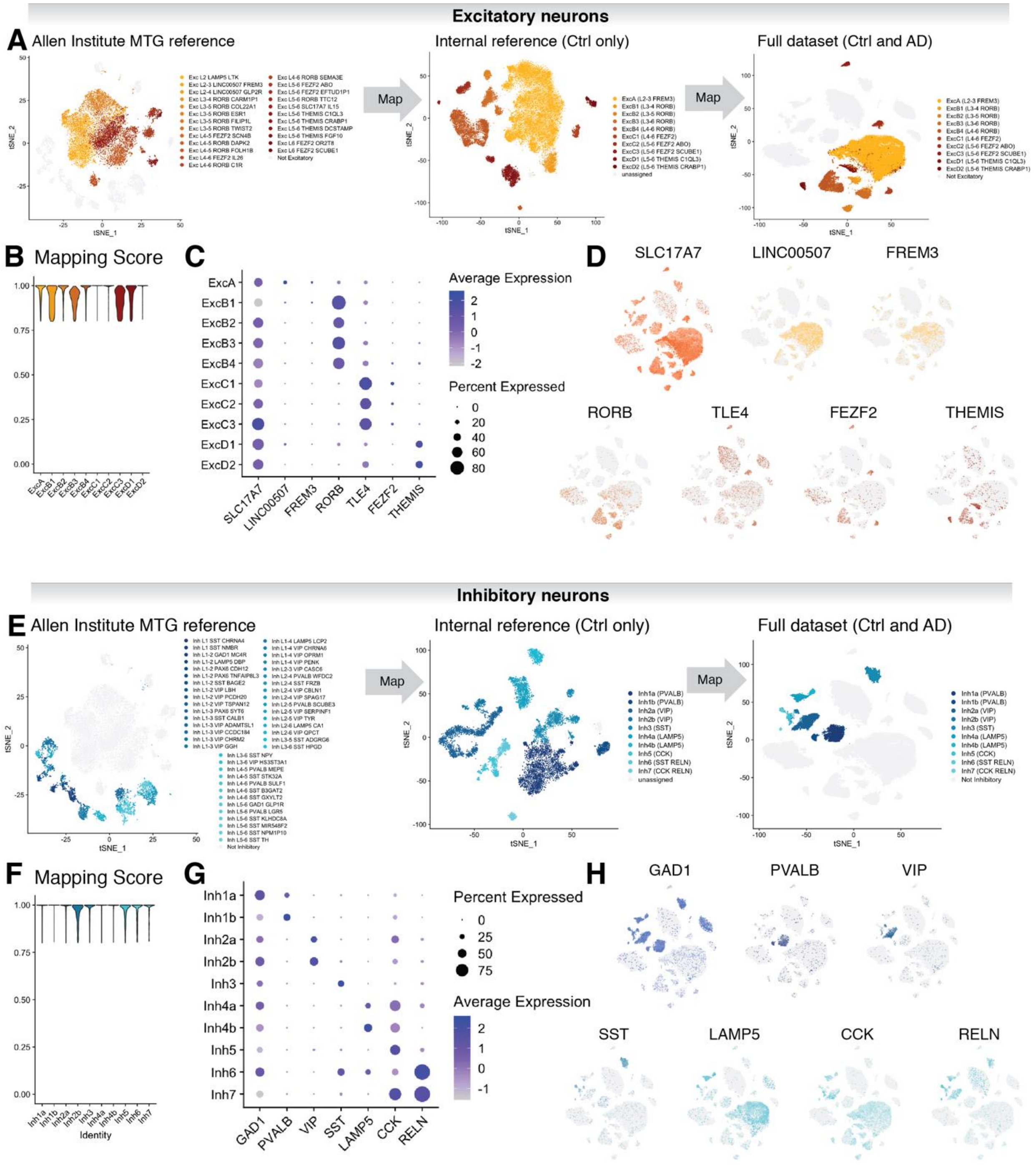


**Supplementary Fig. 4. Mapping of excitatory and inhibitory neuronal subtypes.** (**A, E**) Excitatory (A) or inhibitory (E) neuronal subtypes of the Allen Institute MTG reference were projected onto excitatory (A) or inhibitory (E) neurons of the internal reference and collapsed into 10 subgroups each based on graph-based clustering and mapping scores (see Methods for details). Neuronal subtypes of the internal reference were then projected onto excitatory (A) or inhibitory (E) neurons of the complete dataset. (**B, F**) Violin plots of scores of excitatory (B) or inhibitory (F) cell type mapping of the complete dataset to the internal reference after cleanup. (**C, G**) Dot plot of selected genes distinguishing excitatory (C) or inhibitory (G) neuronal cell types. Size of the dots shows the percentage of nuclei with non-zero expression and color represents the scaled average expression. (**D, H**) Expression of selected genes specific for excitatory (D) or inhibitory (H) neuronal subtypes overlaid on the t-SNE projection of the complete dataset.

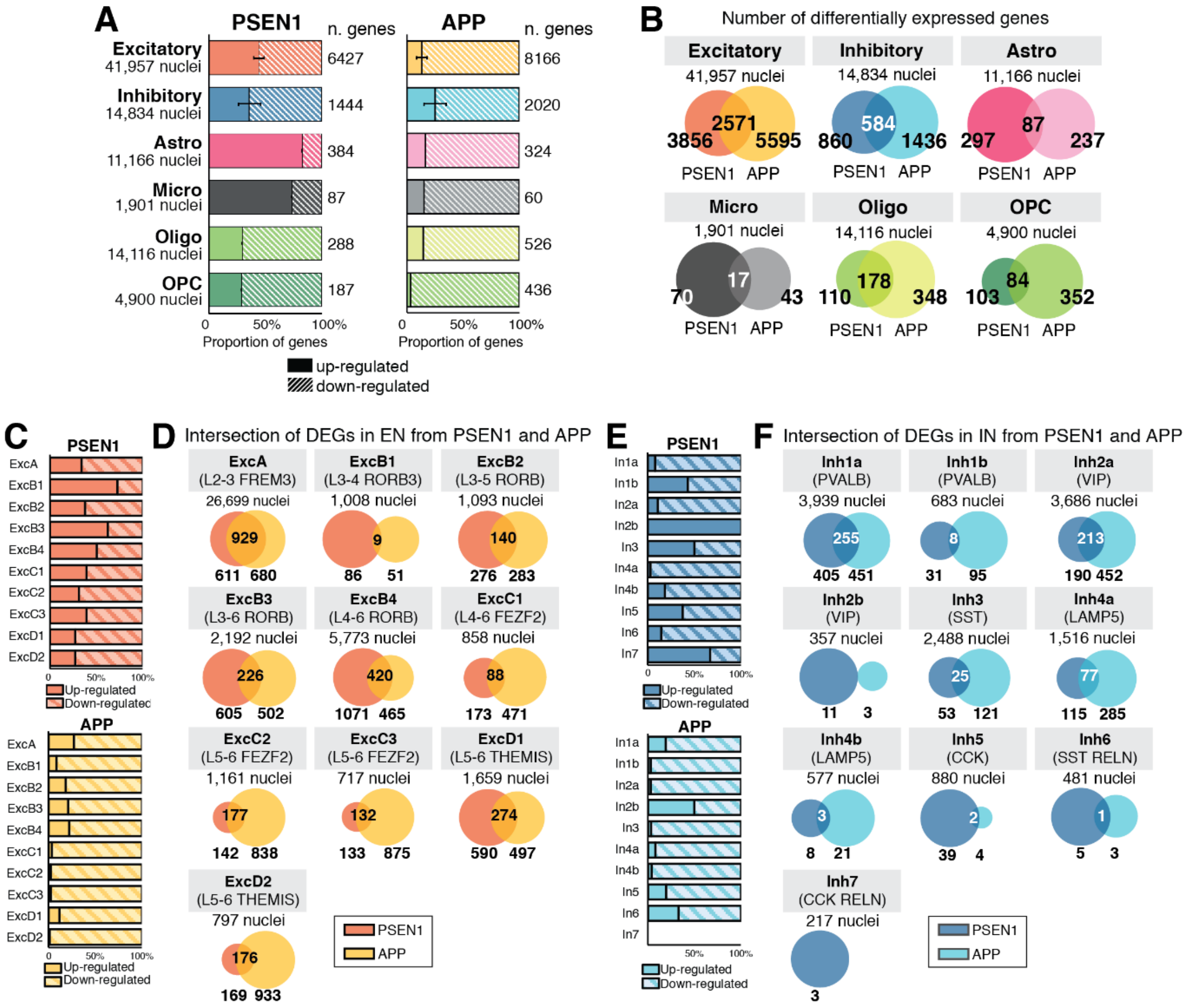


**Supplementary Fig. 5. Differential expression in neurons**

(**A**) Proportion of up- and down-regulated differentially expressed genes in APP and PSEN1 AD samples are shown for each cell type. For excitatory and inhibitory neurons, mean ±SEM of all subtypes is shown; individual values are in (C) and (D). (**B**) Venn diagrams show the number of significantly differentially expressed genes that are detected in each cell type, when comparing cells from APP and PSEN1 AD samples to their respective matched controls. The overlap indicates the number of genes detected differentially expressed in both mutations. (**C-F**) Number of differentially expressed genes in excitatory and inhibitory neuronal subtypes split by mutation. (C, E) Bar graphs showing the percentage of significantly up- and down-regulated genes in excitatory (C) and inhibitory (E) neuronal subtypes. (D, F) Venn diagrams showing the number of differentially expressed genes in APP or PSEN1 AD samples compared to their respective matched non-demented controls in excitatory (D) or inhibitory (F) subtypes.

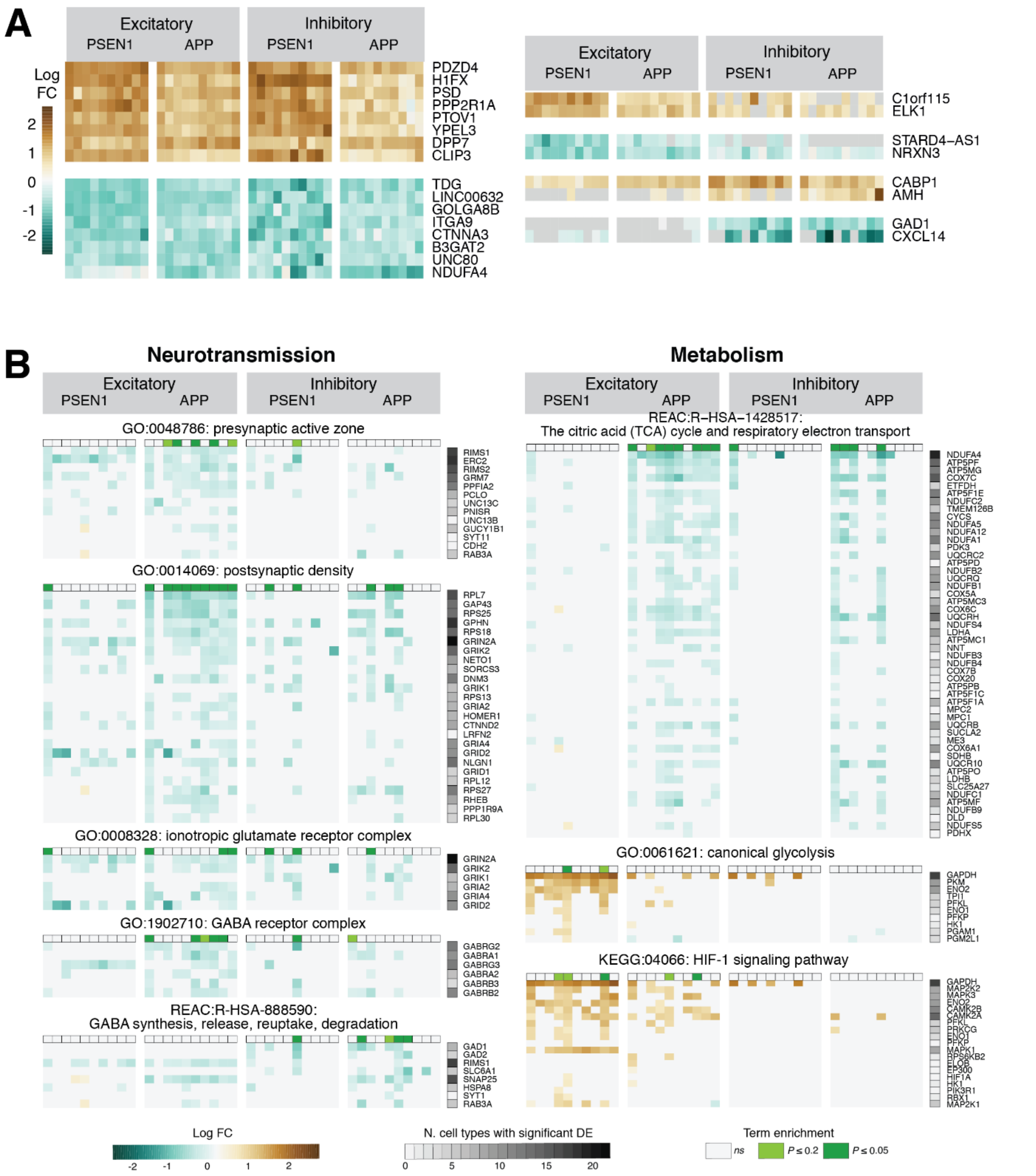


**Supplementary Fig. 6. Differentially expressed genes and altered biological processes in PSEN1 and APP AD neurons**

(**A**) Heatmap of log-transformed fold change of 8 most up- and down-regulated genes across excitatory and inhibitory subtypes, and the 2 most specifically up- and down-regulated genes in excitatory or inhibitory neurons for indicated comparisons of APP or PSEN1 AD samples with the respective matched controls. (**B**) Heatmaps of differentially expressed genes in indicated GO terms and pathways related to neurotransmission and metabolism enriched in up- or down-regulated genes in neurons referred to Fig. 2A, B. Heatmaps show the log-transformed fold change for indicated comparisons of APP or PSEN1 AD samples with their matched controls.

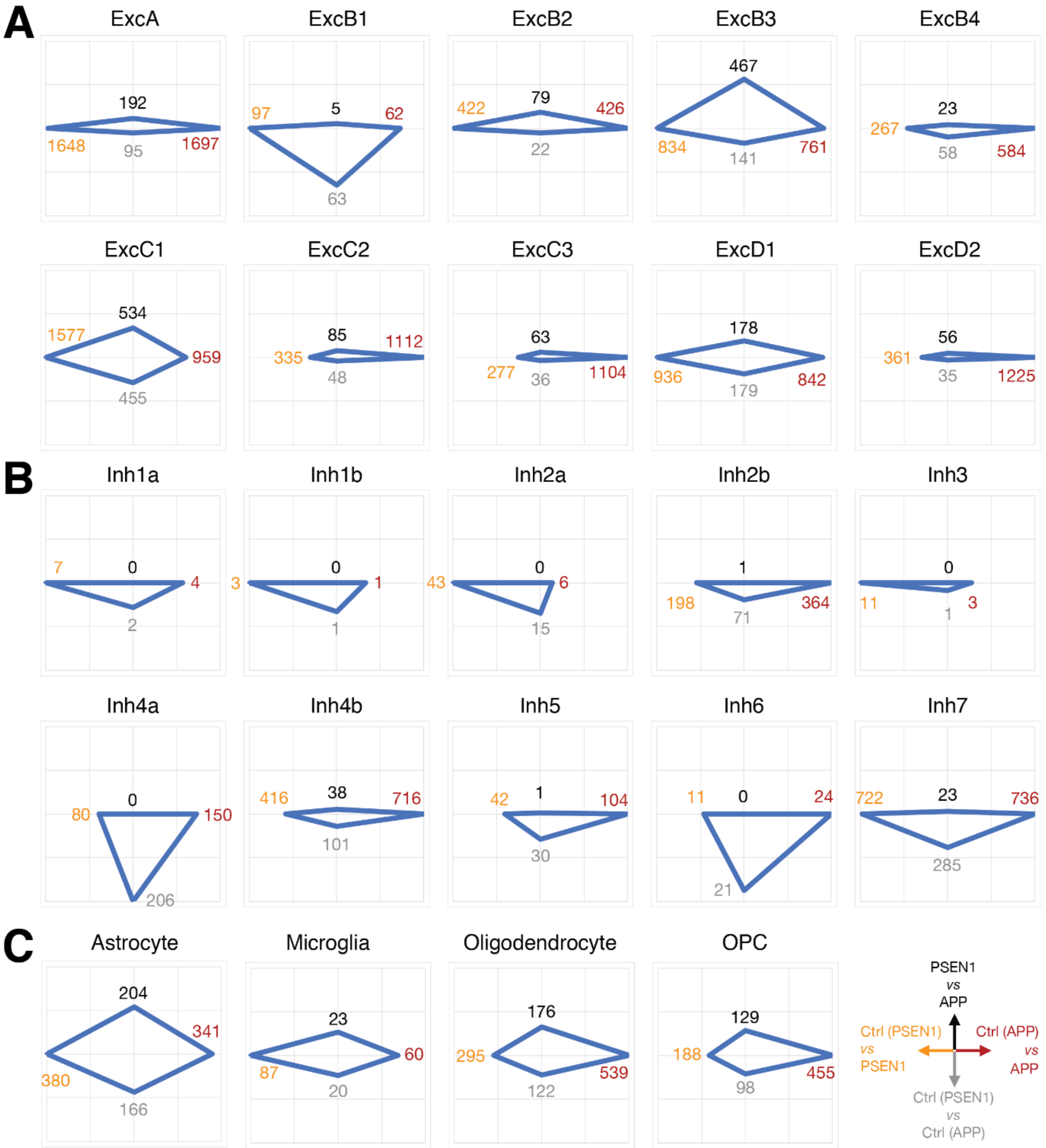


**Supplementary Fig. 7. Number of differentially expressed genes between AD and non-demented controls**

(**A-C**) Number of significantly differentially expressed genes between APP AD and matched controls (left), PSEN1 AD and matched controls (right), APP AD and PSEN1 AD (up) and the two control groups (down) in excitatory (A) and inhibitory (B) neuronal subtypes and glial cell types (C) are plotted, as well as an overview of graph layouts.

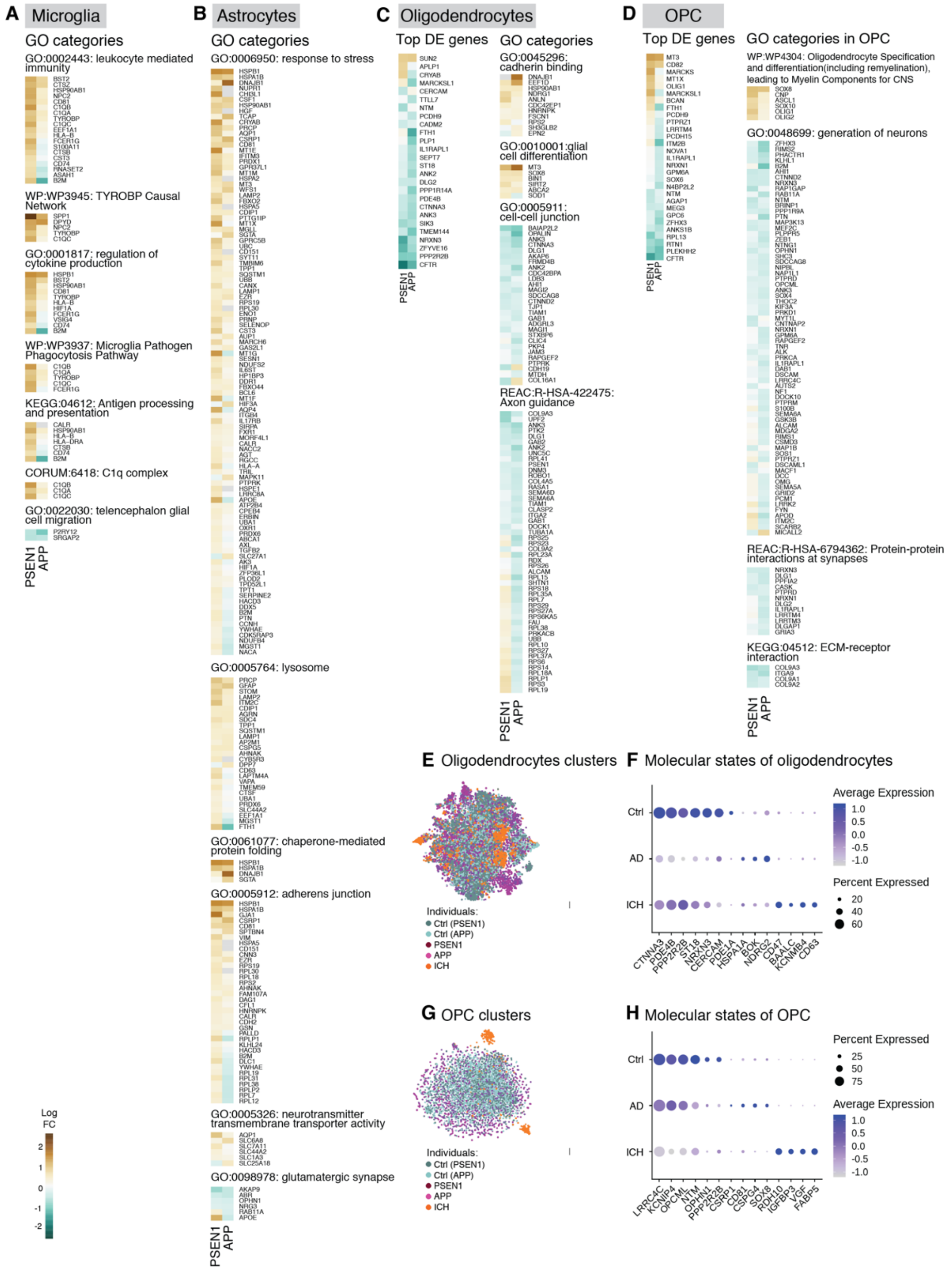

**Supplementary Fig. 8. Altered biological processes and phenotypic changes in glial cells in PSEN1 and APP AD**

(**A-D**) Heatmaps of differentially expressed genes in indicated GO terms and pathways enriched in up- or down-regulated genes in microglia (A; referred to Fig. 3A), astrocytes (B; referred to

Fig. 3B), oligodendrocytes (C) and OPC (D). Heatmaps show the log-transformed fold change for indicated comparisons of APP or PSEN1 AD samples with their matched controls. (**E, G**) Distribution of disease groups in two-dimensional (t-SNE) projections of oligodendrocytes (D) and OPCs (G). (**F, H**) Molecular states of oligodendrocytes (E) and OPCs (H) defined by expression of genes that distinguish controls, AD and intracerebral hemorrhage (ICH). Dot size represents percentage of nuclei with non-zero expression, colors represent scaled average expression.

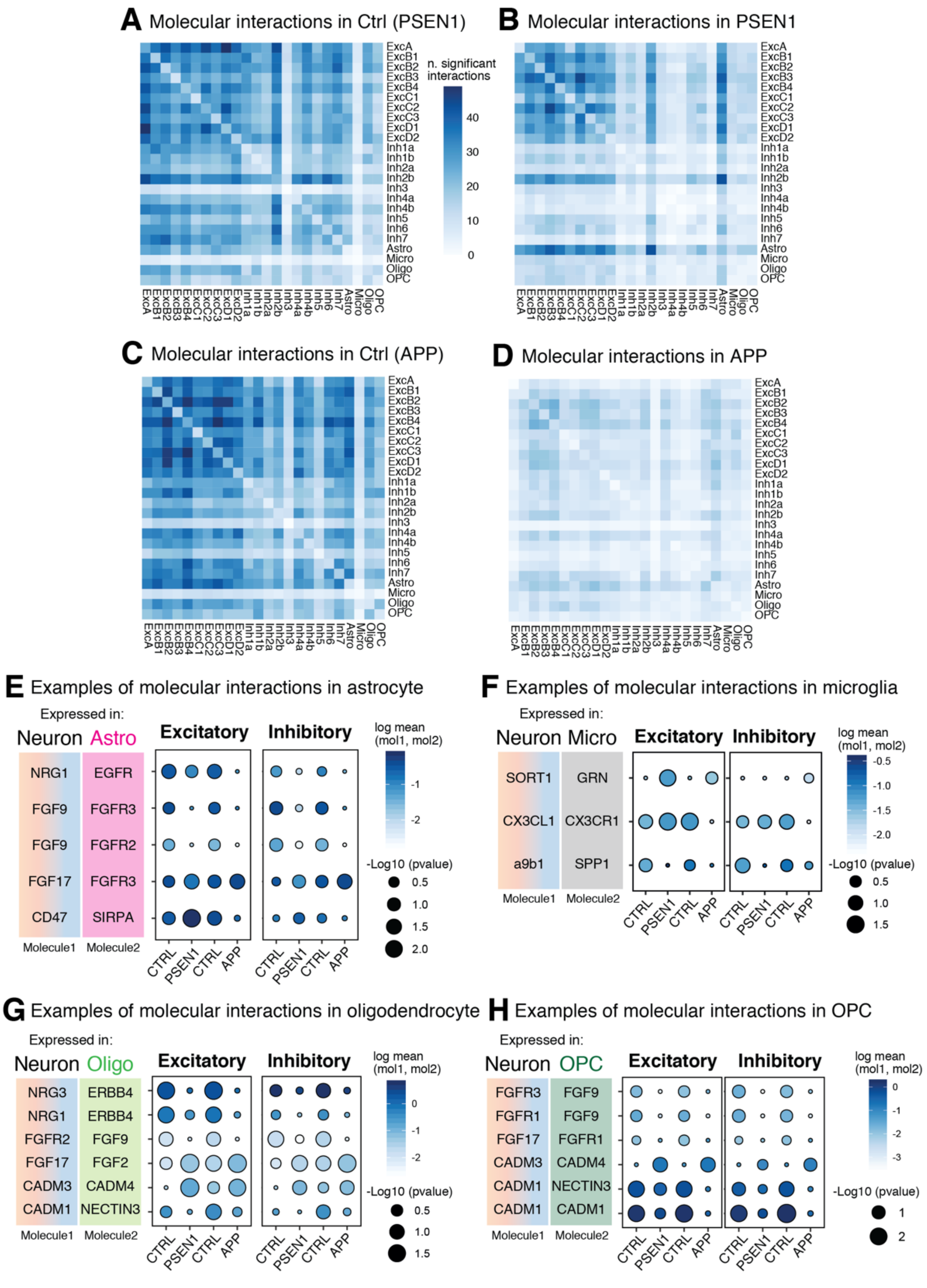


**Supplementary Fig. 9. Analysis of potential molecular interactions between different cell types**

Strength of potential intercellular interactions were analyzed using CellphoneDB (*66*). Heatmaps of the number of significant receptor-ligand pairs identified between indicated cell types in non-demented controls (Ctrl (PSEN1) (**A**), Ctrl (APP) (**C**)), PSEN1 (**B**) and APP (**D**). Examples of molecular interaction pairs in astrocytes (**E**), microglia (**F**), oligodendrocytes (**G**) and OPCs (**H**). Values of average significance score across excitatory or inhibitory neuronal subtypes (circle size) and log-transformed average expression of the interacting molecules in each cell-cell interaction (blue scale) are represented.

**Data S1.**

Sample and clinical information for non-demented controls and Alzheimer's disease patients.

| Patient ID | Age at death | Post mortem delay | Sex | Braak and Braak stage | Mutation |
|---|---|---|---|---|---|
| Ctrl1 | 56 | 82h | Female | - | - |
| Ctrl2 | 59 | 53h | Female | - | - |
| Ctrl3 | 63 | 42h | Male | - | - |
| Ctrl4 | 66 | 81h | Male | - | - |
| Ctrl5 | 45 | 93h | Female | - | - |
| Ctrl6 | 53 | 29.5h | Female | - | - |
| Ctrl7 | 51 | 52h | Male | - | - |
| Ctrl8 | 53 | 96h | Male | - | - |
| AD1 | 56 | 16h | Female | VI | APP V717 |
| AD2 | 59 | 89h | Female | VI | APP V717 |
| AD3 | 62 | 32h | Male | VI | APP V717 |
| AD4 | 66 | 68h | Male | VI | APP V717 |
| AD5 | 41 | 64h | Female | VI | PSEN1 Intron 4 |
| AD6 | 51 | 32h | Female | VI | PSEN1 Intron 4 |
| AD7 | 51 | 43.1h | Male | VI | PSEN1 Intron 4 |
| AD8 | 54 | 115h | Male | VI | PSEN1 M146I |
